# Cholangiocyte primary cilia transduce a fluid shear signal to increase KLF2 via the actin cytoskeleton

**DOI:** 10.1101/2023.03.01.530694

**Authors:** Cody J. Wehrkamp, Andrew M. Oleksijew, Adrian P. Mansini, Sergio A. Gradilone, Ashley M. Mohr, Justin L. Mott

## Abstract

Cholangiocarcinoma is an aggressive solid tumor formed in the bile duct epithelium. Often this tumor obstructs bile flow, known as cholestasis. Normal cholangiocytes detect bile flow in the ductal lumen with an extension of the apical membrane called the primary cilium. However, these sensory organelles are often lost in malignant cells. Krüppel-like factor 2 (KLF2) is an important flow-sensitive transcription factor involved in shear stress response in endothelial cells, and has anti-proliferative, anti-inflammatory, and anti-angiogenic effects. The potential role of KLF2 in cholangiocyte flow detection and in cholangiocarcinoma is unknown. We hypothesized that reduced bile flow contributes to malignant features in cholangiocarcinoma through regulation of KLF2 signaling. We observed that primary cilia were expressed in normal cholangiocytes but were absent in malignant cells. KLF2 expression was higher in normal cells compared to malignant. Depletion of cilia in normal cells led to a decrease in KLF2 expression and increased cilia number was associated with increased KLF2. Enforced KLF2 expression inhibited cell proliferation, migration, and also decreased cell death induction in malignant cells. Applied media flow over cholangiocytes increased KLF2 and cilia depletion completely blocked flow-induced KLF2 expression. Disruption of filamentous actin decreased KLF2 expression, suggesting the cilium may communicate through a cytoskeletal mechanotransduction pathway. Our studies demonstrate that cilia positively regulated KLF2 protein levels and increased fluid flow induced KLF2 expression for the first time in cholangiocytes, emphasizing the importance of reestablishing bile flow in cholestatic cholangiocarcinoma.

## Introduction

Cholangiocarcinoma is an aggressive epithelial neoplasm of the biliary tree associated with inflammation, injury, and bile duct obstruction. This cancer is often diagnosed at a late stage due to its silent character, and unresectable tumors are uniformly lethal. The five-year survival is less than 15%, and intrahepatic cholangiocarcinoma is the most aggressive subtype with a five-year survival of 6%.^1^ The asymptomatic nature of this disease portends a reduced opportunity for surgical resection, with only approximately 35% of patients considered eligible for surgery with curative intent.^2^ Absent resection, the current standard of care is chemotherapy; however, cholangiocarcinoma is characteristically chemoresistant and these therapies are palliative only.^3,4^ Cholangiocarcinoma tumors are particularly heterogeneous, due to the diversity of anatomical location, varied predisposing risk factors, and wide assortment of associated genetic alteration.^5^ These features contribute to a narrow range of treatment options, and at the same time, create a need for expansion of understanding of the specific molecular pathways exploited by this cancer.

Biliary obstruction is the most common cause of the presenting symptoms of cholangiocarcinoma.^6^ Bile duct obstruction can arise either from conditions that predispose to cancer, such as primary sclerosing cholangitis (PSC) or intrahepatic cholelithiasis, or from tumor-induced ductal blockage. Premalignant inflammation (PSC or liver fluke infestation) can limit bile flow through strictures that narrow the lumen.^7,8^ Intrahepatic bile stones, while not a common cause, can induce both obstructive cholestasis and cholangiocarcinoma.^9,10^ The tumor itself can decrease or block flow through compression of the bile duct by mass expansion effect. To recapitulate this feature of human disease, several of the existing cholangiocarcinoma animal models induce cholestasis with bile duct ligation.^11^ Moreover, the cholestatic environment in combination with chemical carcinogen treatment in some models is required for carcinogenesis.^12,13^ Additionally, in an orthotopic or oncogenic transduction model, induced cholestasis by bile duct ligation significantly increased tumor progression.^14,15^ In the healthy biliary system, cholangiocytes sense and maintain the regular flow of bile. Reduced flow can induce cholangiocyte proliferation contributing to some cholangiopathies including polycystic liver disease.^16^ The mechanisms by which cholestasis promotes cholangiocarcinoma are not fully defined.

The primary cilium has been well documented as an important signaling organelle in many cell types in the human body.^17,18^ Functioning as a sensory antenna for the cell, the primary cilium is responsive to mechanical, chemical, and osmotic stimuli. Its significance in various biological processes is revealed by the pathologies to which its dysfunction or absence can be associated, or directly attributed. Often, cancer cells lack or have defective primary cilia compared to their non-malignant counterparts.^19–22^ This is also true for biliary epithelial cells. Normal cholangiocytes possess primary cilia, with which they can detect bile flow in the ductal lumen; however, primary cilia are lost in cholangiocarcinoma cells.^23,24^

Krüppel-like factor 2 (KLF2) is a member of a large family of transcription factors that direct a range of cellular processes including proliferation, apoptosis, differentiation, inflammation, and migration.^25–29^ It is not surprising then that the expression or function of many KLFs is dysregulated in cancer. ^26,30,31^ Separately, considerable study has established KLF2 as a key factor involved in shear stress response to flow in non-malignant endothelial cells, where under normal, undisturbed flow conditions, KLF2 expression exerts anti-proliferative and anti-inflammatory protective effects.^32,33^ The status and function of KLF2 in cholangiocarcinoma is not yet known. Furthermore, the capacity of KLF2 as a mediator of flow response in a non-endothelial cell type has not been described.

Here, our studies have demonstrated that KLF2 may be an important mediator of cholangiocyte behavior in settings of healthy bile flow and absent in cholestasis. KLF2 was both decreased in cholangiocarcinoma cells lines as well as patient cholangiocarcinoma tissue. KLF2 was found to respond to shear stress in cholangiocytes dependent on the primary cilium-actin cytoskeletal axis, demonstrating a molecular mechanism for healthy mechanosensation of bile flow in cholangiocytes. Cholangiocarcinoma cell lines were found to have decreased cilium expression, with decreased KLF2 levels, but overexpression of this transcription factor decreased migration and proliferation in these cell lines. Our data support KLF2 as a mediator of the bile flow response of cholangiocytes in the bile duct and in cholestasis its loss contributes to aggressive phenotypes in cholangiocarcinoma.

## Methods

### Cell lines

Human cholangiocarcinoma cell lines were previously derived from a female patient with metastatic gallbladder cancer, Mz-ChA-1 [289], a male patient with combined histologic features of hepatocellular carcinoma and cholangiocarcinoma, KMCH [290], and from the malignant cells of ascites from a male patient with intrahepatic cholangiocarcinoma, HuCCT-1. ^34–36^ Recently-derived patient cholangiocarcinoma cell lines ICC2, ICC3, ICC8, and ICC11 were from resected intrahepatic cholangiocarcinoma tumors.^37^ H69 cells are an SV-40 transformed non-malignant immortalized cholangiocyte cell line.^38^ NHC cells are normal human cholangiocytes isolated from normal tissue during surgical dissection of a local hepatic adenoma in a female patient.^39^ Human umbilical vein endothelial cells (HUVECs) were purchased from American Type Culture Collection (ATCC CRL-1730). Cholangiocarcinoma cells were grown in Dulbecco’s modified Eagle medium (DMEM) supplemented with 10% fetal bovine serum (FBS), insulin (0.5 µg/mL) and G418 (50 µg/mL). Recently-derived patient cell lines were grown in Roswell Park Memorial Institute (RPMI) medium supplemented with 20% FBS, 1% L-glutamine, 1% MEM non-essential amino acids solution, 1% sodium pyruvate, 50 µg/mL G418, and 5 µg/mL insulin on collagen-coated culture dishes. H69 and NHC cells were grown in DMEM supplemented with 10% FBS, insulin (5 µg/mL), adenine (24.3 µg/mL), epinephrine (1 µg/mL), T3-T (triiodothyronine (T3)-transferrin (T), [T3-2.23 ng/ml, T-8.19 µg/mL]), epidermal growth factor (9.9 ng/mL) and hydrocortisone (5.34 µg/mL). HUVECs were grown in F-12K medium with 10% FBS, 0.1 mg/mL heparin, and 5 mL endothelial cell growth supplement (ECGS, BD Biosciences).

Cholangiocarcinoma is a rare cancer and there is no central source of cholangiocarcinoma cell lines to be used as standards for validation by genetic testing. We employ immunostaining for keratin-19, an intermediate filament protein and marker of cholangiocyte differentiation to indicate that a cell is cholangiocyte derived. Additionally, all members of the lab are trained to identify the unique cell morphologies of the cholangiocyte cells when grown in standard culture conditions and to monitor cell appearance with each experiment.

### Tissue isolation

Frozen samples were obtained from patient-resected normal liver or cholangiocarcinoma tumor tissues acquired from the UNMC Tissue Bank. Samples were kept frozen until isolation and all materials for isolation were sterile or bleached and rinsed, and then chilled with liquid nitrogen or dry ice before contact with tissues. Tissues were transferred from dry ice to a mortar containing a small volume of liquid nitrogen and ground to a fine powder with a chilled pestel.

Samples were lysed in RNA lysis buffer (mirVana, see RNA transfection methods), or protein lysis buffer (see Immunoblotting methods), and immediately frozen in a dry ice/isopropanol bath or used for isolation by the traditional protocols described.

### Quantitative reverse-transcription PCR

Total RNA for mRNA qRT-PCR was isolated using TRIzol Reagent (Life Technologies). Two micrograms of RNA were reverse-transcribed using Moloney leukemia virus reverse transcriptase and random hexamers. Quantitative real-time PCR was performed using SYBR Green DNA binding (Roche). Fifteen nanograms of RNA was converted to cDNA using specific primers for KLF2 (Forward: ACTCACACCTGCAGCTACGC; Reverse: GCACAGATGGCACTGGAAT) or 18S (Forward: CGTTCTTAGTTGGTGGAGCG, Reverse: CGCTGAGCCAGTCAGTGTAG) (Applied Biosystems). qRT-PCR was performed using RedTaq master mix (Applied Biosystems) and specific TaqMan primers (Applied Biosystems) on an iCycler (BioRad) thermocycler. Relative RNA expression was calculated using the delta CT method.

### Immunoblotting

Cells were lysed in lysis buffer containing 50 mM Tris-HCl (pH 7.4), 150 mM sodium chloride, 1 mM EDTA, 1 mM DTT (dithiothreitol), 1 mM sodium orthovanadate, 100 mM sodium fluoride, and 1% Triton X-100 (w/v) supplemented with complete protease inhibitors (Roche). Lysates were incubated on ice for 15 minutes and vortexed at 5-minute intervals. After lysis, insoluble proteins were removed by centrifugation at 15,700 *g* for 10 minutes. Soluble protein was resolved by sodium dodecyl sulfate-polyacrylamide gel electrophoresis (SDS-PAGE), transferred to nitrocellulose. The membrane was blocked with 5% dry milk (w/v) in TBST for 45-60 minutes and incubated with the indicated primary antibody for KLF2 (Aviva Systems Biology ARP32760), IFT88 (proteintech 13967-1-AP), or actin (Sigma A2228) overnight at 4°C. Incubation with HRP-conjugated secondary in blocking buffer was performed at room temperature for 1 hour. Protein bands were detected using enhanced chemiluminescence (ECL) and radiographic film.

### Immunofluorescence

Cells were seeded onto glass coverslips that were pre-coated with collagen by application of collagen solution (Sigma, Cat # C8919) with a cotton swab and allowing it to dry. After drying, coverslips were added to culture dishes containing growth medium and cells plated as usual. The coverslips were transferred to a new 6-well plate and rinsed with PBS at room temperature. They were fixed for 20 minutes at 37 °C in PBS with 3% paraformaldehyde, 100 mM piperazine-1,4-bis-2-ethanesulfonic acid (PIPES), 3 mM MgSO_4_, and 1 mM EGTA. After washing 3 times with PBS, the cells were permeabilized for 5 minutes with 0.1% Tween-20 in PBS. Cells were washed again 3 times in PBS then blocked for 60 minutes at 37 °C in blocking solution containing PBS with 5% glycerol, 5% donkey serum, and 0.0 % sodium azide. Primary antibodies were diluted 1:200-1:500 in blocking solution and 150 µL was pipetted on top of each coverslip. Arl13-b antibody (proteintech 17711-1-A) was incubated overnight at 4 °C in a 6-well plate that was sealed with parafilm to prevent evaporation. The coverslips were transferred to a new 6-well plate and washed 3 times with PBS. They were then incubated with 1:2000 secondary Alexa Fluor antibodies (Alexa488, Alexa594, Thermo Fisher) in blocking buffer for 45-60 minutes at 37 °C. After washing 3 times in PBS and with a final water rinse to remove salts, coverslips were mounted onto glass slides in a drop of Vectashield with DAPI (Vector Laboratories) mounting medium. Immunofluorescence images were captured using a Zeiss 710 confocal laser scanning microscope in the UNMC Advanced Microscopy Core Facility.

### Nuclear morphology apoptosis assay

Treated living cells were stained with the nuclear dye 4’6-diamidino-2-phenylindole (DAPI; Sigma, Cat # D9542) at a final concentration of 5 µg/mL for 20 minutes at 37 °C prior to live cell imaging by epifluorescence (Leica DMI6000B). Cells were counted as DAPI-positive if the nucleus showed bright staining and as apoptotic if there was characteristic nuclear fragmentation, blebbing, or pyknosis. Total cell number was determined in the same field by phase contrast microscopy, and data are expressed as the percent of DAPI-positive nuclei out of the total.

### Caspase 3/7 activity assay

Cells were seeded in a 96-well plate in 200 µL medium. The following day, they were treated or transfected for 24-48 hours. Cell death was induced with treatment of 4-50 ng/mL TRAIL or 5 μg/mL staurosporine for 6 hours. Then, 170 µL medium was removed and 30 µL caspase 3/7 reaction (1:100 Z-DEVD-R110 fluorogenic substrate:Homogenous buffer; ApoONE, Promega) was added to each well. The plate was covered with aluminum foil and mixed by gentle shaking for at least 30 seconds, then allowed to incubate for 1-16 hours. Fluorescence was measured at 535 nm on a Tecan Infinite 200 Pro plate reader. The pan-caspase inhibitor Z-VAD-fmk (Santa Cruz, Cat # sc-3067) was resuspended in DMSO and used at a final working concentration of 50 µM.

### MTT proliferation assay

Cell proliferation was assayed by reduction of 3-(4,5-dimethylthiazol-2-yl)-2,5-diphenyltetrazolium bromide (MTT; Invitrogen, M6494). MTT was freshly dissolved into PBS at a stock concentration of 12 mM and diluted into phenol red-free DMEM with 10% FBS for a final MTT concentration of 2 mM. Reactions were carried out at 37 °C for four hours and stopped by removing the medium. Reduced MTT was dissolved in 100 µL isopropanol and absorbance measured at 540 nm on a plate reader. All data are corrected to the initial signal, set at 100%. Assays were repeated four times for each condition.

### Transwell assay

Twenty-five thousand cells in 100 μL serum-free medium were seeded into the upper inserts of a Corning Costar 24-well 8-μm membrane pore size plate containing 600 μL regular medium with 10% FBS in the bottom of the wells. Cells were allowed to migrate for 24 to 72 hours. To mount the membranes for quantification of migrated cells, inserts were rinsed with PBS, then the upper side of the membrane was thoroughly wiped with two cotton swabs to remove adhering cells that had not migrated. Cells were fixed by dipping the insert 20 times in isopropanol followed by a water rinse. The membrane was removed from the insert with a scalpel and mounted cells-down on a glass slide with a drop of ProLong Gold antifade reagent with DAPI (Thermo Fisher) and covered with a cover slip. Cells were imaged by epifluorescence on a Leica DMI6000B microscope.

### RNA transfection

siRNA against IFT88 (ID: 149319, Ambion) or Negative Control siRNA (Cat# 1027310, Qiagen) were reverse-transfected using Lipofectamine RNAiMAX (Thermo Fisher) to a final oligonucleotide concentration of 20 nM by seeding cells directly onto the transfection reaction.

### Stable transfection

Cells were transfected with a KLF2 expression plasmid containing a puromycin resistance marker and a DsRed expression cassette (pBRPyCAG-hKlf2-DsRed-Ip; addgene, Cat # 26277) using Lipofectamine 3000 (Thermo Scientific). Clones were selected by treatment with 2 μg/mL puromycin in normal medium that was replaced every few days. Puromycin-resistant cells were reseeded thinly in selection medium to allow for isolated growth of colonies from individual cells. Colonies were screened for positive DsRed fluorescence and transferred to new dishes. Positive clones were then further screened for KLF2 expression by qRT-PCR and immunoblot.

### Orbital shear stress model

To induce shear stress over a cell monolayer, cells were seeded in barrier-separated peripheries (as below, in blue) of 10-cm culture dishes and rotated on an orbital shaker. To create each of these “racetracks,” a 6-cm dish was fixed in the center of a 10-cm dish with a small volume of chloroform to fuse the plastic together (diagram shown in **Figure 3A)**.The fused dishes were set aside to allow any remaining chloroform to dissipate before cells were seeded.

Cells were seeded in 8 mL of medium in the peripheral annulus, and shear stress was generated with rotation at 80 or 120 rpm for 24 to 96 hours on a remote CO_2_-resistant orbital shaker (Thermo Scientific, Cat # 88881101) inside a cell incubator under normal culture conditions. Shear stress was calculated using the formula 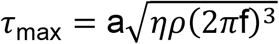 where a is the radius of rotation defined by the rotary plate (0.85 cm), *η* is the viscosity of the medium (0.0101 poise), *ρ* is the density of the medium (1.007 g/mL), and f is the frequency of rotation (rotations/second).^40–42^ This resulted in shear stress values within the physiologic range of the force of bile flow on the wall of the bile duct.^43,44^

### Statistical analysis

Data were analyzed by ANOVA with *post hoc* Bonferroni correction when multiple comparisons were possible. When only two conditions were measured, 2-tailed, equal variance Student’s *t-*test was employed. Comparisons between groups were considered significantly different when the p-value was less than or equal to 0.05. Where applicable: * = p < 0.05, ** = p < 0.01, *** = p < 0.001.

## Results

### Expression of KLF2 was decreased in cholangiocarcinoma

We considered that cilia may regulate the transcription factor KLF2 and examined the expression of KLF2 in normal cholangiocyte cells and malignant cholangiocarcinoma cells. KLF2 protein expression was higher in normal H69 cells compared to cholangiocarcinoma cell lines HuCCT and KMCH (**Figure 1A**), as well as OZ and EGI-1 (not shown). We expanded our analysis of KLF2 expression status in malignant cells by sampling recently derived cell lines from four cholangiocarcinoma patients. These cells, ICC2, ICC3, ICC8, and ICC11, exhibited lower KLF2 protein expression compared to normal NHC cells (**Figure 1B**). Additionally, human tissue samples were as follows: three normal livers, three samples of normal liver adjacent to cholangiocarcinoma, and 14 cholangiocarcinoma tumor samples. We probed for KLF2 protein by immunoblot and actin as a control. We quantified the relative KLF2 protein signal versus actin and calculated the mean relative KLF2 expression. KLF2 expression in tumor samples was significantly lower than in healthy controls. We also observed a decrease KLF expression in the tumor-adjacent normal liver tissue, but this did not reach statistical significance (**Figure 1C**). Together, these data demonstrate a consistent decrease in KLF2 expression in cholangiocarcinoma.

**Figure 1.**
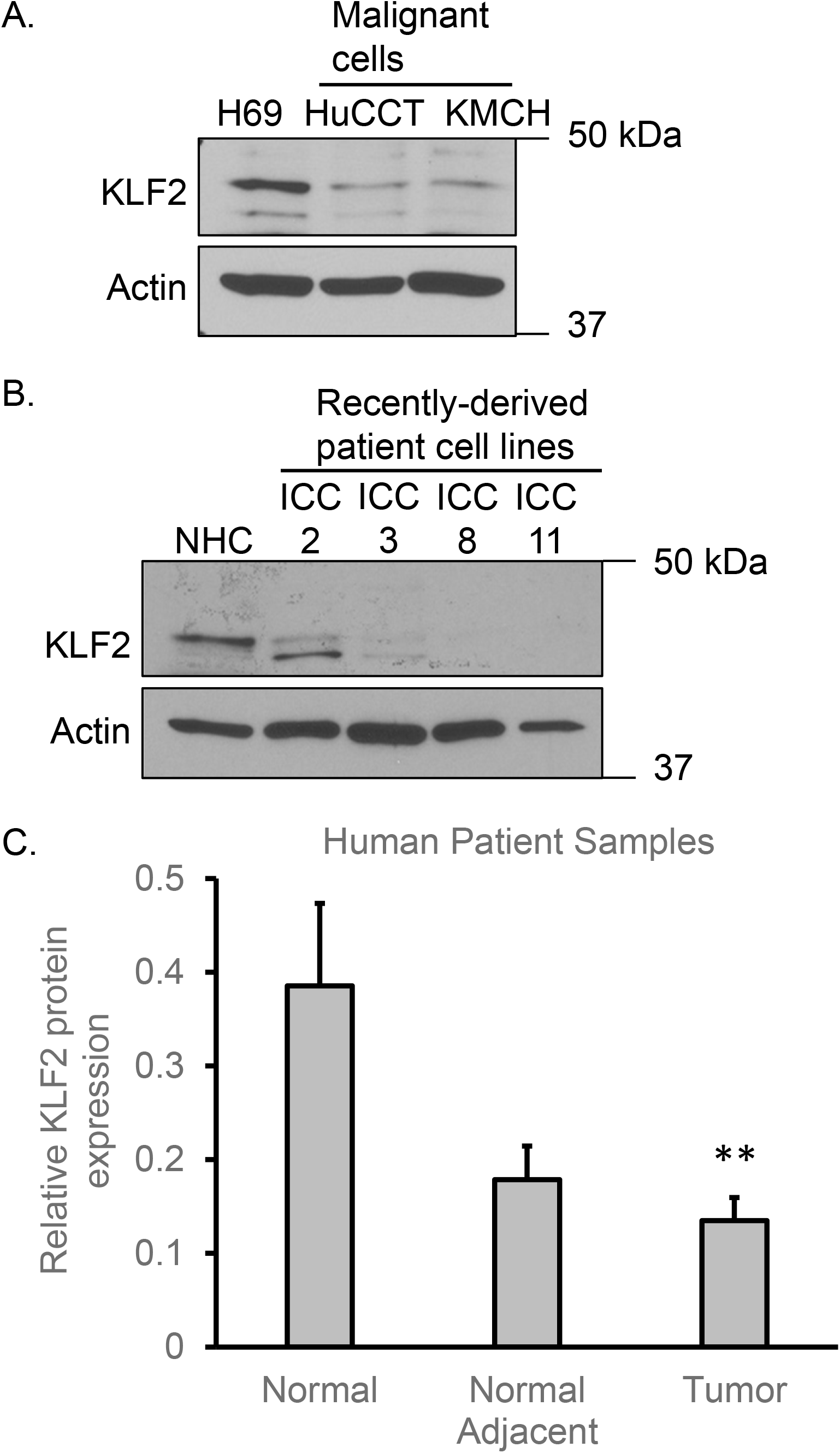
Expression of KLF2 is decreased in cholangiocarcinoma. Normal cholangiocyte cell line H69 has higher KLF2 protein expression than cholangiocarcinoma cell lines HuCCT and KMCH (**A**). Patient-derived cholangiocarcinoma cells have decreased KLF2 expression compared to normal cholangiocyte cell line NHC (**B**). **Panel C**: Relative KLF2 protein expression compared to actin loading control was measured by signal on immunoblot. Normal human liver tissue expressed significantly higher KLF2 protein than cholangiocarcinoma tumors. ** p < 0.01, using ANOVA with *post hoc* correction.

### Primary cilia and KLF2 expression in cholangiocytes were correlated

We sought to determine the presence of primary cilia in cultured cholangiocyte and cholangiocarcinoma cell lines. Our previous reports have shown a reduced frequency or absence of cilia in malignant cells compared to non-malignant cells.^45^ Serum deprivation was demonstrated to induce ciliogenesis and is a commonly used practice to promote cilia formation *in vitro*.^45,46^ We cultured non-malignant cholangiocyte cell lines, NHC and H69, and malignant cholangiocarcinoma cells, Mz-ChA-1, KMCH, and HuCCT, in serum-free medium for 24 hours. Cells were then assessed by fluorescence microscopy for presence of cilia. Specifically, we detected ADP-ribosylation factor-like protein 13B (Arl13b), a cilia-localized protein that assists in cilia formation and maintenance.^47^ Imaging revealed significant expression of cilia in normal cholangiocyte cells (**Figure 2A & B**), while cholangiocarcinoma cells expressed essentially none (**Figure 2C-E**). These data demonstrate that malignant cholangiocarcinoma cells lack primary cilia.

**Figure 2.**
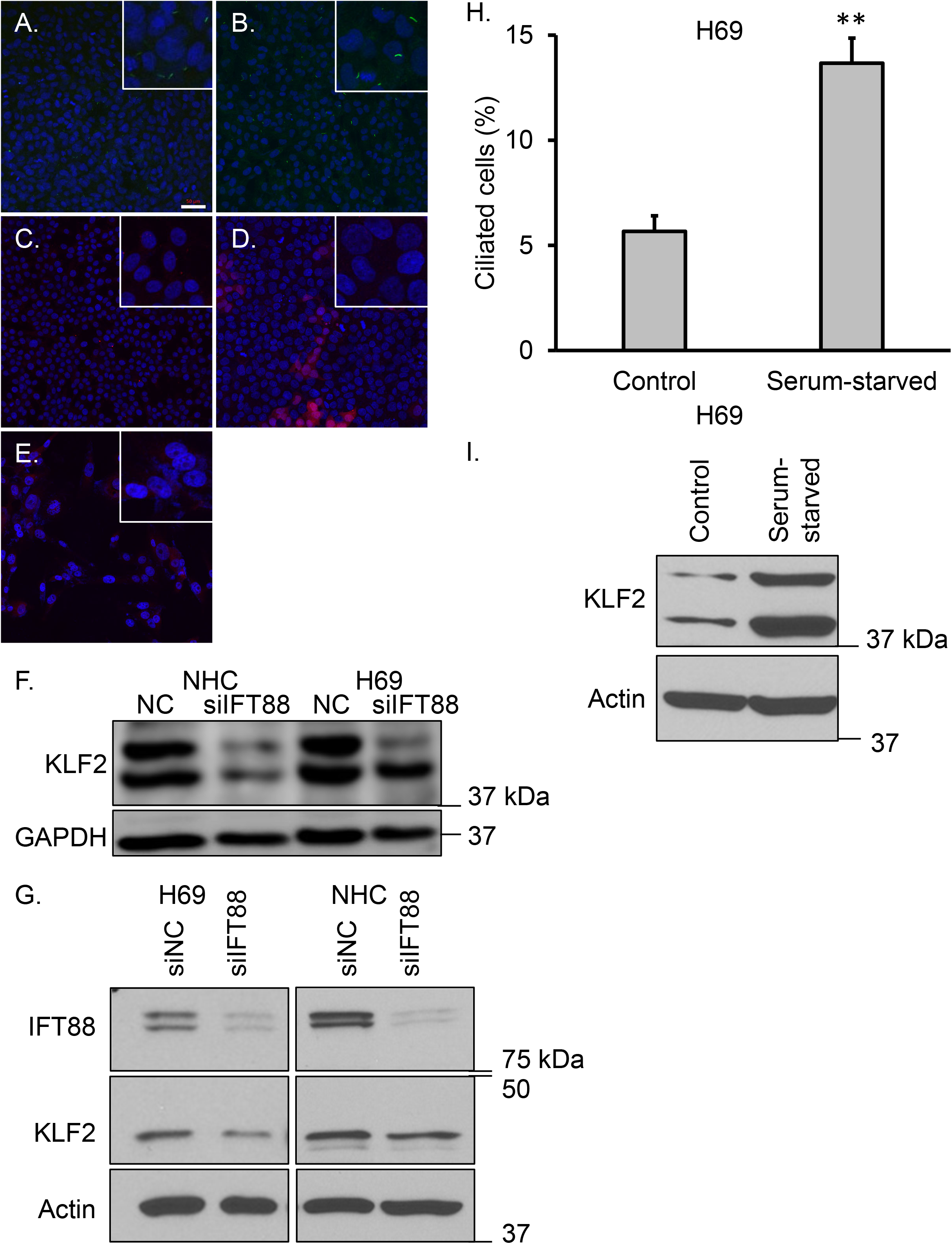
Normal but not malignant cholangiocytes express primary cilia. Upper panels. Normal cholangiocyte cells NHC (**A**) and H69 (**B**) were fixed and probed for the ciliary marker Arl13b (green) and visualized by confocal fluorescence microscopy. Lower panels. Cholangiocarcinoma cells Mz-ChA-1 (**C**), KMCH (**D**), and HuCCT (**E**) were probed with anti-Arl13b (red). Cilia-expressing NHC and H69 cells had higher KLF2 expression than cells with cilia knocked down by treatment with siRNA against IFT88, a protein required for ciliogenesis and maintenance (**F**). IFT88 protein was knocked down by siRNA in cholangiocytes and KLF2 protein expression decreased (**G**). Normal cholangiocytes were serum-starved for 48 hours, which led to an increase in percentage of ciliated cells (**H**), and an increase in KLF2 expression compared to cells grown in serum-containing medium (**I**). Nuclei were stained with DAPI. Scale bar = 50 μm. All images are at the same scale. Inserts: each insert is a 3-fold magnification of a subsection of the respective image. Quantification of ciliated cells by confocal microscopy is based on the average of three separate high-powered fields. Actin or GAPDH were probed as loading controls. ** p < 0.01, Student’s *t*-test.

To investigate the expression pattern and potential link between the cholangiocyte primary cilium and KLF2, we experimentally ablated cilia in cholangiocyte cells. This was accomplished with RNAi suppression of intraflagellar transport 88 (IFT88), a ciliary transport protein necessary for the biogenesis and maintenance of primary cilia.^23,48^ Our previous data showed that knockout of IFT88 in cholangiocyte cell lines H69 and NHC resulted in complete loss of primary cilia expression.^23,46^ De-ciliation of NHC and H69 cells caused a decrease in KLF2 protein (**Figures 2F-G**). An established means of promoting ciliogenesis in epithelial cells is by serum deprivation (starvation). This is based on the reciprocal link between ciliogenesis and the cell cycle.^49–51^ Serum starvation for 48 hours resulted in a significant increase in the percentage of ciliated cholangiocytes compared to cells grown in serum-containing medium (**Figure 2H**). Serum-starved cells also showed a robust increase in KLF2 protein compared to controls (**Figure 2I**). These results indicate a positive correlation between the presentation of primary cilia and the expression of KLF2 in cholangiocytes and treatments to increase or decrease cilia caused increased or decreased KLF2.

### Fluid flow induced KLF2 in cholangiocytes

Because KLF2 is known to be responsive to shear stress generated by fluid flow in endothelial cells, we wanted to simulate this effect and test if it is a flow-responsive factor in cholangiocytes. First, we tested our orbital flow model (**Figure 3A)** in the human endothelial cell line HUVEC and observed a robust increase in KLF2 mRNA expression with the application of arterial levels of shear stress over 48 hours (**Figure 3B**). Cholangiocyte epithelial cells do not experience this higher shear, therefore, the level of shear stress was reduced to physiologic bile flow levels. Cholangiocyte cells responded similarly as endothelial cells, and flow induced a significant increase in KLF2 mRNA compared to static controls (**Figure 3C-D**). This is the first demonstration of KLF2 regulation by fluid flow in cholangiocytes, to our knowledge.

**Figure 3.**
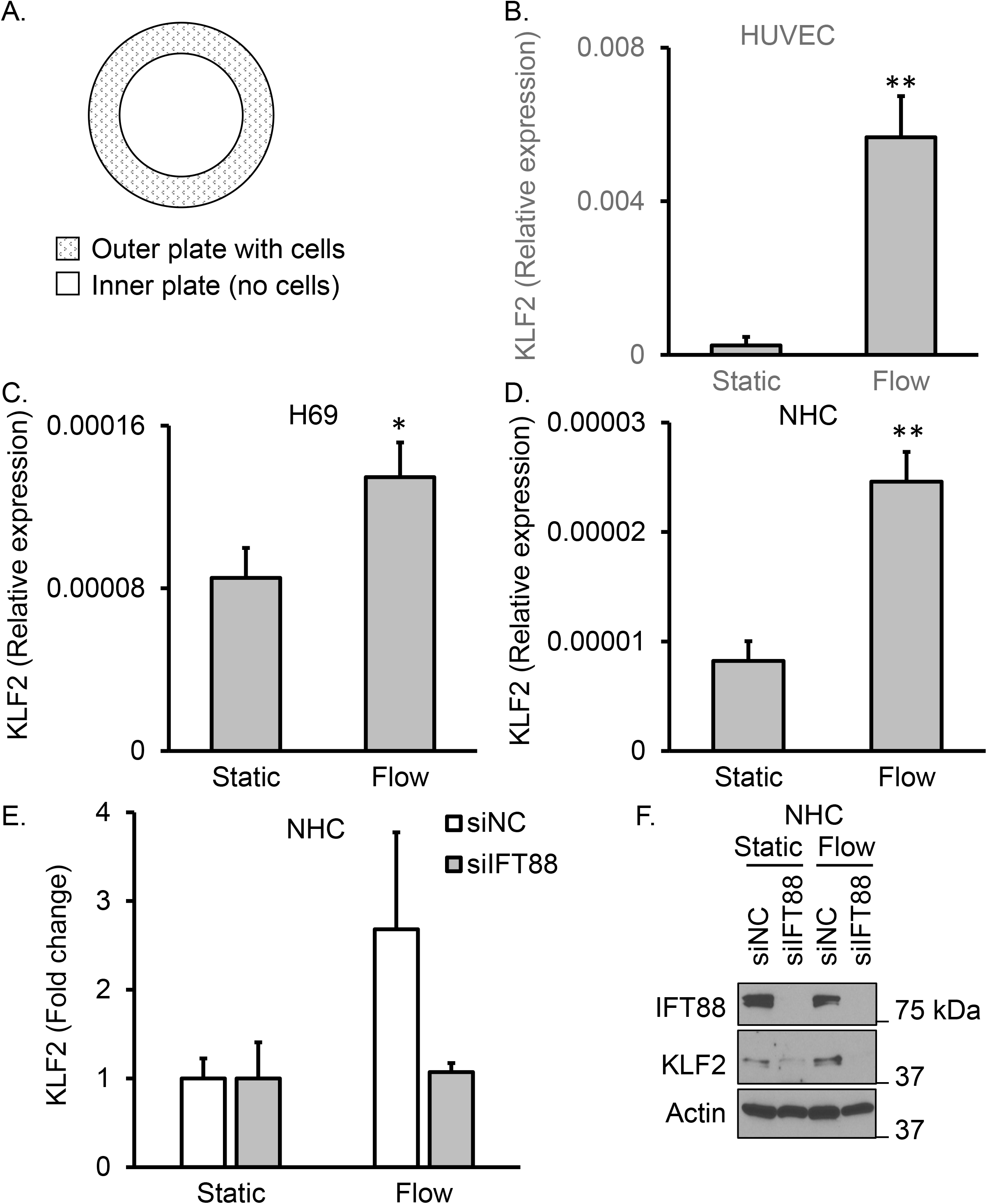
Flow-induced KLF2 expression is cilia-dependent in cholangiocytes. Cells were subjected to physiologic shear stress and compared to cells with no flow (**A**). Fluid shear stress induced a significant increase in KLF2 mRNA in endothelial cells (HUVEC) (**B**), and cholangiocytes (**C**,**D**), as measured by qRT-PCR. NHC cells were transfected with control siRNA (siNC) or siIFT88 to remove cilia and then incubated under static or flow conditions. Fold-change of KLF2 mRNA increased by flow with siNC but this induction was abolished with siIFT88 transfection. 18S rRNA was used as a control RNA and actin as a protein loading control (**E**). Similarly, knockdown of IFT88 protein decreased KLF2 protein under static conditions, and IFT88 knockdown prevented induction of KLF2 protein expression by flow (**F**). * p < 0.05, ** p < 0.01; using ANOVA with *post hoc* correction.

Fluid flow can induce opening of mechanosensitive ion channels, including the cilium-localized PC2 calcium channel.^6^ To investigate if calcium is acting as a second messenger in the flow-induced signal transduction of KLF2, we used the cell-permeant calcium chelating molecule 1,2-bis(2-aminophenoxy)ethane-N,N,N′,N′-tetraacetic acid tetrakis(acetoxymethyl ester) (BAPTA-AM) to scavenge cytosolic calcium and again exposed cells to fluid flow. There was no significant difference in KLF2 induction when calcium was chelated (not shown). This suggests that the KLF2 flow response is not significantly dependent on calcium in cholangiocytes, and that another signaling mechanism is likely at play.

To further characterize the KLF2 flow response in cholangiocytes, we evaluated the primary cilium as a candidate flow detector. Toward this end, we again employed siRNA knockdown of IFT88 to generate de-ciliated cells and then measured KLF2 expression under static or flow conditions. NHC cells were transfected with control siRNA (siNC) or siRNA to IFT88 (siIFT88) for 16 hours and then incubated under static or flow conditions for an additional 48 hours. Shear stress caused an increased fold-change of KLF2 mRNA in siNC-transfected cells; however, this increase was abolished in cells with cilia ablation by siRNA to IFT88 (**Figure 3E**). Fluid flow in siNC control cells induced KLF2 protein expression. In NHC cells under static conditions, removal of cilia with siRNA to IFT88 led to a drop in KLF2 compared to control.

Moreover, IFT88 knockdown prevented flow induction of KLF2 protein expression (**Figure 3F**). The KLF2 flow response could be completely abolished by removal of primary cilia, placing KLF2 downstream of primary cilia signaling and revealing a ciliary-dependent mechanism of KLF2 flow response in cholangiocyte cells.

### Actin disruption uncoupled primary cilia and KLF2 expression

The ciliary axoneme communicates with the actin cytoskeleton, and depolymerization of cytoplasmic actin can induce increased cilia formation.^52,53^ We postulated that the increase in cilia induced by the actin-depolymerizing small molecule cytochalasin D would cause an increase in KLF2 as well. We treated H69 cells with vehicle (DMSO) or 100 μM cytochalasin D for 24 hours, then probed for cilia-specific marker Arl13b and visualized by fluorescence confocal microscopy. Vehicle-treated cells displayed typical morphology and cilia expression (**Figure 4A**). Treatment with cytochalasin D caused a change in cell shape, likely due to disruption of the actin cytoskeleton, and also caused an increase in the number of cells expressing cilia, as well as a qualitative increase in the length of cilia in these cells (**Figure 4B**). Quantification of cilia-expressing cells compared to the total cell number showed a significant increase in cilia expression with actin filament disruption (**Figure 4C**). Under the same conditions to promote actin cytoskeleton destabilization, instead of increased KLF2 there was an unexpected decrease in KLF2 protein (**Figure 4D**) and mRNA levels (**Figure 4E**). Thus, disruption of actin filaments uncoupled the signal from cilia to increase KLF2.

**Figure 4.**
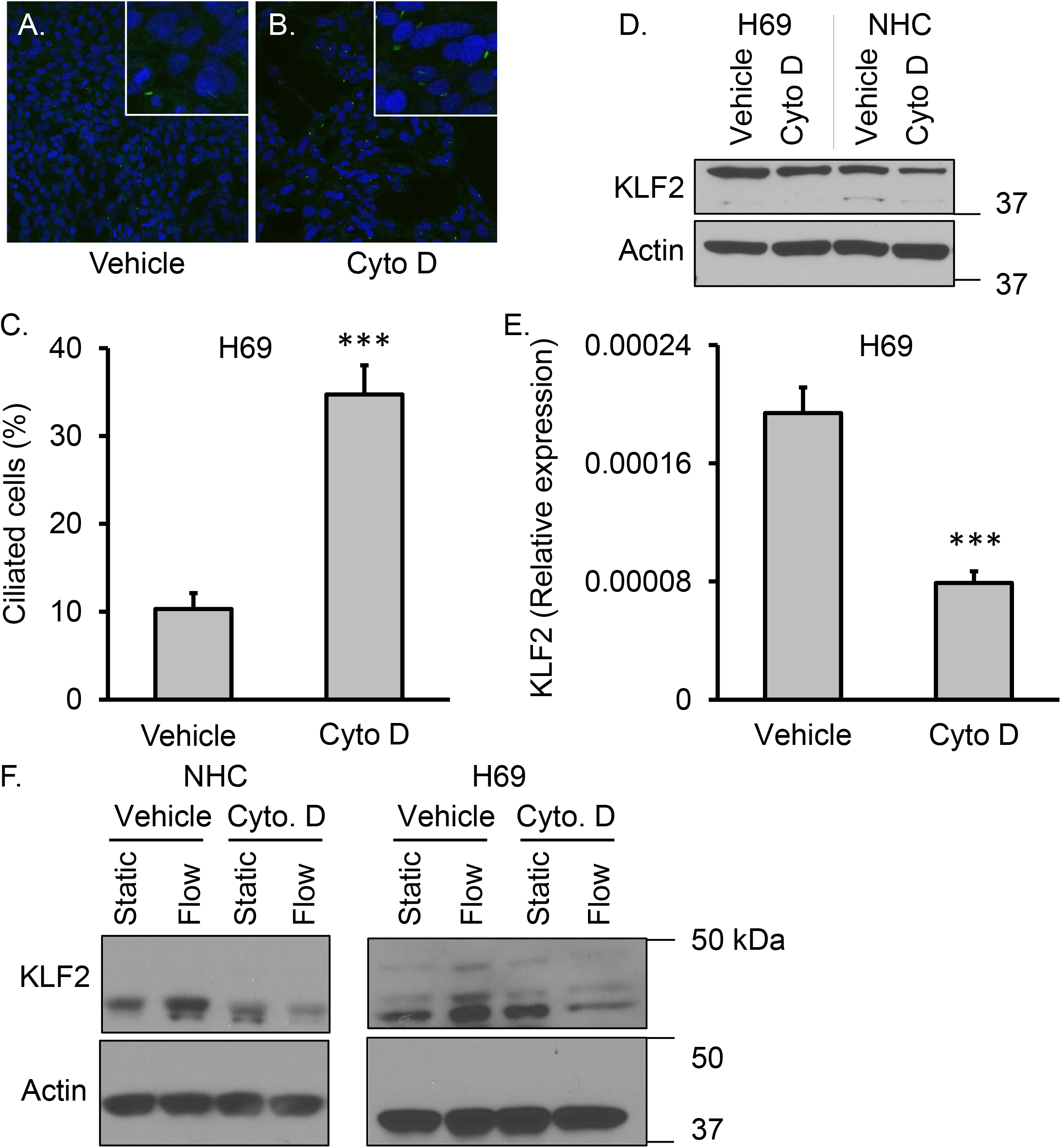
Flow-induced KLF2 expression is dependent on intact actin cytoskeleton. Normal cholangiocytes were treated with vehicle (**A**) or 100 μM cytochalasin D (Cyto D), an inhibitor of filamentous actin polymerization (**B**), for 24 hours and primary cilia were imaged by confocal fluorescence microscopy.The percentage of ciliated cells was increased by cytochalasin D treatment (**C**). Actin filament disruption caused a decrease in KLF2 protein expression (**D**) and mRNA expression (**E**). 18S rRNA was used as a control RNA and actin as a protein loading control (**E**). Increased expression of KLF2 in response to flow was abolished by cytochalasin D in NHC and H69 cholangiocytes (**F)**. *** p < 0.001.

As KLF2 expression decreased with disruption of the actin cytoskeleton, we hypothesized that cholangiocyte mechanosensation of shear stress through KLF2 depends on intact actin cytoskeleton. To assess the potential of the actin cytoskeleton to act as a downstream signal of the KLF2 fluid shear response, NHC and H69 cholangiocytes were subjected to static or bile physiologic shear conditions (flow), with or without 100 μM of cytochalasin D. Disruption of the actin cytoskeleton blocked the increase in KLF2 protein levels that was seen in the shear stress conditions (**Figure 4E-F)**. This demonstrates the importance of intact actin cytoskeleton in the mechanosensory response of this transcription factor.

### KLF2 overexpression decreases proliferation, migration, and apoptosis

The functional implications of KLF2 regulation were determined by directly manipulating KLF2 expression. Cells were transfected with a KLF2 expression construct that contained a puromycin resistance cassette and dsRed fluorescent marker for selection of stable clones.

Selected clonal KMCH cells and HEK293T cells stably overexpressed KLF2 at the mRNA (**Figure 5A & B**) and protein levels (**Figure 5C**). KLF2-overexpressing cells proliferated at a slower rate that became significant beginning at 24 hours for KMCH (**Figure 5D**) and 48 hours for HEK293T (**Figure 5E**) and continued through 72 hours.

**Figure 5.**
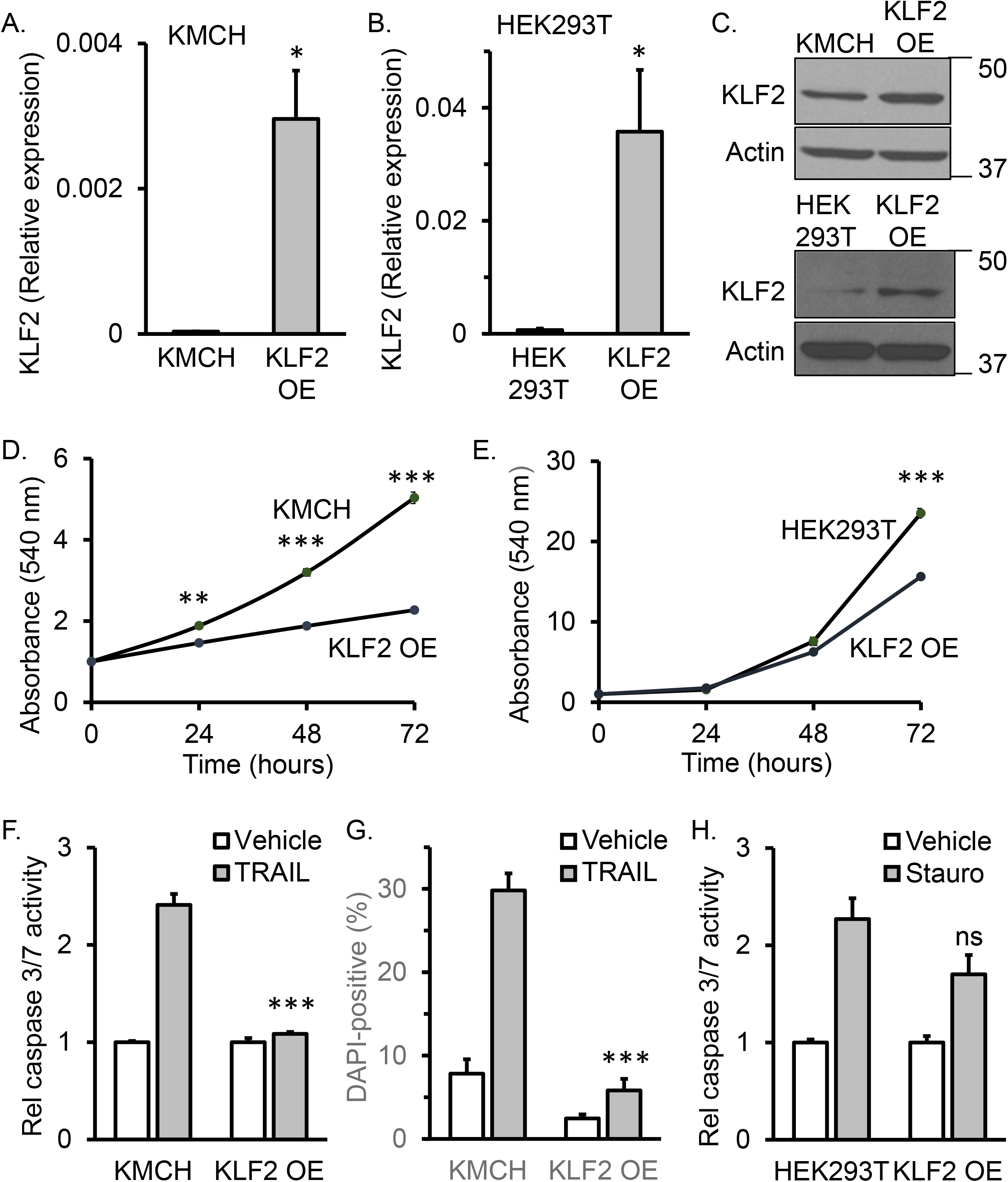
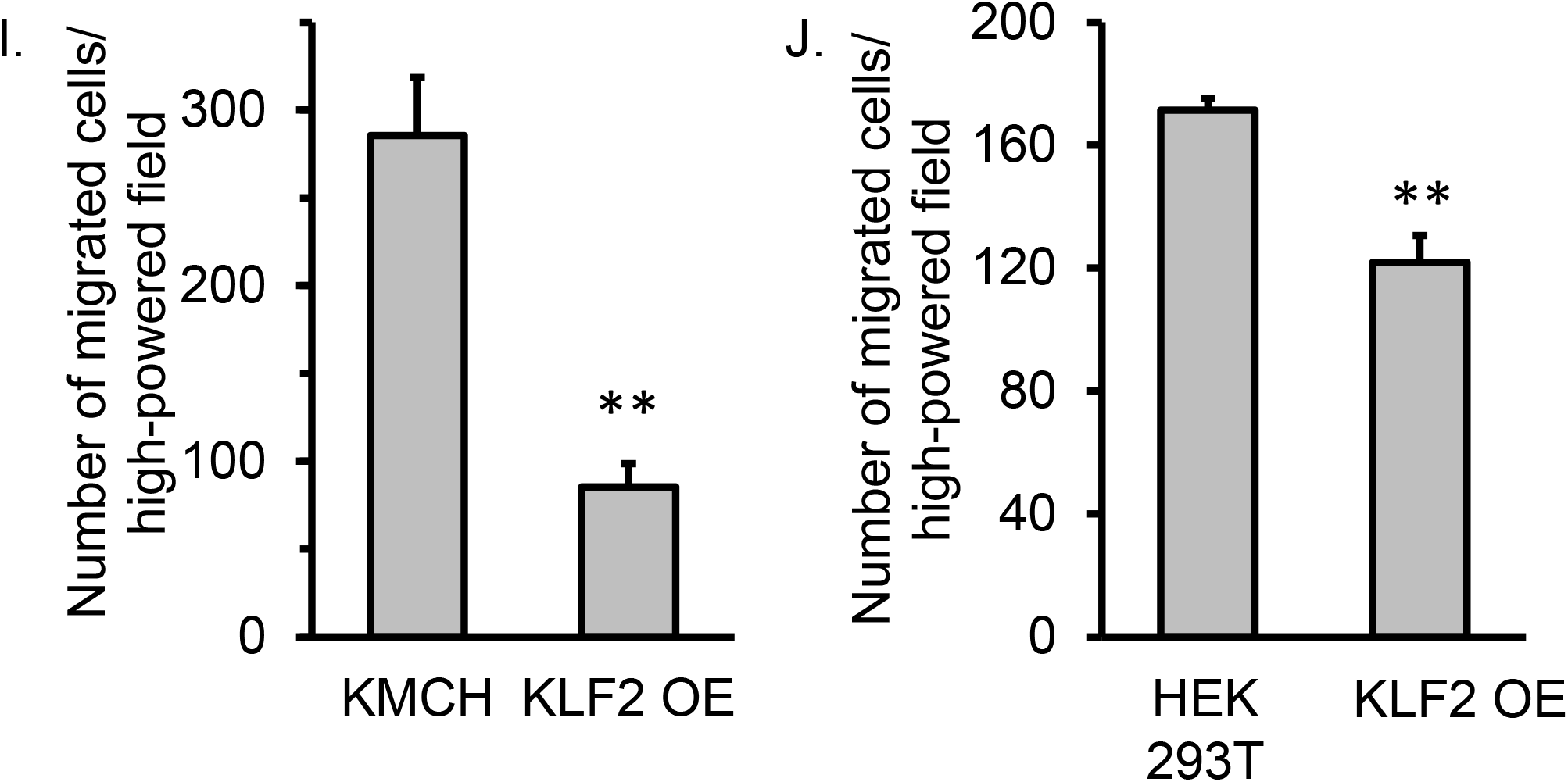
KLF2 overexpression decreased proliferation, apoptosis, and migration in cholangiocarcinoma cells. Stable transfection with a KLF2 expression plasmid of KMCH (**A**) or HEK293T (**B**) cells generated overexpression (OE) clones with increased KLF2 as measured by qRT-PCR (**A & B**) or immunoblot (**C)**. Proliferation was significantly decreased in KLF2 overexpression (OE) clones compared to parental KMCH (**D**) and HEK293T (**E**) cells as measured by MTT assay over 72 hours. KLF2-normal or -overexpressing KMCH cells were treated with vehicle or 50 ng/mL TRAIL to induce cell death and HEK293T cells were treated with vehicle or 5 μg/mL staurosporine. TRAIL-induced apoptosis was decreased in KLF2-overexpressing cells when measured by caspase 3/7 activity assay (**F**) or DAPI staining and nuclear morphology (**G**). KLF2 overexpression in HEK293T cells did not significantly affect staurosporine-induced apoptosis compared to KLF2-normal cells as measured by caspase 3/7 activity (**H**). Cell migratory potential was measured using a transwell assay. Fewer KLF2-expression cells migrated compared to KLF2-normal cells over 72 hours for KMCH (**I**) or 48 hours for HEK293T (**J**). Quantification of data are a mean of three experiments +/-SEM. * p < 0.05, ** p < 0.01, *** p < 0.001, ns = not significant; using Student’s t-test or ANOVA with *post hoc* correction.18S rRNA was used as a control RNA for qRT-PCR, and actin as a loading control for western blot.

We tested the effect of KLF2 overexpression on resistance to apoptosis. In KMCH cells, treatment with TRAIL to induce cell death revealed decreased sensitivity in KLF2-overexpressing cells versus control. This effect on apoptosis was observed when measured by activation of effector caspases 3 and 7 (**Figure 5F**) or by nuclear morphology changes seen with retention and condensation of DAPI stain (**Figure 5G**). HEK293T cells were found to be TRAIL-insensitive, possibly because they are not malignant cells, thus staurosporine was employed to induce apoptosis. Induction of cell death in HEK293T cells resulted in an increase in relative caspase activity for both control and KLF2-overexpressing cells that was not significantly different (**Figure 5H**). These data indicate that KLF2 caused reduced apoptosis in cholangiocarcinoma cells, but may not have the same apoptosis-suppressing effect in non-malignant cells. Alternatively, cell line-specific effects may be unrelated to the malignant status, a possibility that remains to be determined. The experiments in Figure 5 included the HEK293T cell line because it is readily transfected and we could not generate a second cholangiocyte-derived cell line that stably overexpressed KLF2, a limitation of these data.

We tested the role of KLF2 in cell migratory activity using a transwell assay. Identical numbers of cells were seeded in serum-free medium in the upper section of a Boyden chamber separated by a porous membrane from the lower chamber. The lower chamber contained FBS in the medium as a chemoattractant. Cells that migrated through the membrane were fixed and stained with DAPI and counted by fluorescence microscopy. There was a significant suppression of migratory potential in KLF2-overexpressing clones compared to control cells (**Figures 5I & J**). Overall, enforced expression of KLF2 in malignant cholangiocarcinoma cells decreased proliferation, apoptosis, and migration, consistent with a more quiescent state.

## Discussion

We demonstrated that cilia and KLF2 showed positive correlation and depleting cilia demonstrated this relationship was causal. Our data demonstrated that the KLF2 increase in response to fluid shear stress was dependent on the cholangiocyte primary cilium. The intermediary messaging remains incompletely described, however. Calcium was a potential candidate as it has been shown to be a second messenger in kidney cell ciliary signaling. Fluid shear stress caused calcium-induced calcium release (CICR) via the cilia-localized mechanosensor PC1 and cation channel PC2.^54^ Luminal flow activated these same receptors on the cholangiocyte cilium to increase intracellular calcium as well.^55^ Disruption of calcium signaling inhibited flow induction of KLF2 mRNA in endothelial cells.^56^ Furthermore, the purinergic receptor P2X4, an ATP-driven calcium channel, could mediate KLF2 response to shear stress.^57^ Surprisingly, modulation of intracellular calcium levels by chelation in cholangiocytes did not result in a repression of KLF2 induction by shear stress. Under fluid flow conditions, the increased KLF2 expression compared to static controls was not significantly different with or without calcium chelation. Studies to date that describe calcium-mediated KLF2 signaling have all been performed in endothelial cells. It is possible that the KLF2 signaling axis in epithelial cells—or even biliary cells specifically—employs different biochemical messengers.

In pursuit of other candidate signaling intermediates, we considered the actin cytoskeleton based on reports of its role in primary cilium assembly/disassembly and signal transduction.^53,58,59^ When we manipulated the expression of cilia by preventing actin polymerization, we observed an expected increase in number of cilia. We observed an unexpected decrease in KLF2 expression however. The unexpected direction of change in KLF2 expression is consistent with actin as an intermediate messenger in the primary cilium-KLF2 signaling axis. In summary, in cholangiocytes shear stress and cilia increased KLF2 expression. Bile flow and cilia are decreased in cholangiocarcinoma and we found KLF2 was also decreased in tumors. KLF2 regulated tumor-related features of migration, proliferation, and cell death. Thus, the cholangiocyte primary cilium transduces a fluid shear signal to increase KLF2 and loss of cilia in malignancy reduces KLF and promotes malignant behaviors.

## Acknowledgements

Supported by the National Cancer Institute (NCI) of the National Institutes of Health, award number R01CA222649 (JLM) and 5R01CA183764 (SAG). Additional support from the Fred & Pamela Buffett Cancer Center. CJW was supported by a UNMC Graduate Student Fellowship. We thank the University of Nebraska Medical Center Advanced Microscopy Core Facility, which receives support from the National Institute for General Medical Science (NIGMS) INBRE - P20GM103427 and COBRE - P30GM106397, as well as support from the NCI for The Fred & Pamela Buffett Cancer Center-P30CA036727, and the Nebraska Research Initiative. This publication’s contents and interpretations are the sole responsibility of the authors and do not necessarily represent the official views of the NIH.

## Abbreviations

(BAPTA-AM): 1,2-bis(2-aminophenoxy)ethane-N,N,N′,N′-tetraacetic acid tetrakis(acetoxymethyl ester)
(MTT): 3-(4,5-dimethylthiazol-2-yl)-2,5-diphenyltetrazolium bromide
(DAPI): 4’6-diamidino-2-phenylindole
(ATCC): American Type Culture Collection
(CICR): Calcium-induced calcium release
(dithiothreitol): DTT
(DMEM): Dulbecco’s modified Eagle medium
(ECGS): Endothelial cell growth supplement
(ECL): Enhanced chemiluminescence
(FBS): Fetal bovine serum
(HUVECs): Human umbilical vein endothelial cells
(KLF2): Krüppel-like factor 2
(PIPES): Piperazine-1,4-bis-2-ethanesulfonic acid
(PSC): Primary sclerosing cholangitis
(RPMI): Roswell Park Memorial Institute
(SDS-PAGE): Sodium dodecyl sulfate-polyacrylamide gel electrophoresis

